# Sphingolipid remodelling in SPT-related neuropathies

**DOI:** 10.64898/2026.04.01.715856

**Authors:** Nicole Ziak, Thorsten Hornemann, Museer A. Lone

## Abstract

Sphingolipid homeostasis is critical for neuronal structural and functional integrity in the central and peripheral nervous systems. The rate-limiting enzyme of this pathway, serine-palmitoyltransferase (SPT), establishes the metabolic entry point into sphingolipid biosynthesis. Mutations in the SPT subunits, SPTLC1 and SPTLC2 lead to contrasting disease phenotypes in patients, including amyotrophic lateral sclerosis (ALS) and hereditary sensory neuropathy (HSAN1). A third mixed sensory-motor phenotype is attributed to distinct mutation sets in SPTLC1 and SPTLC2. However, a direct comparison of the metabolic consequences of mutations spanning these disease conditions has not been performed.

Here, we demonstrate that SPTLC1- and SPTLC2-ALS variants contribute to enhanced sphingolipid flux while the ceramide-mediated homeostatic control is impaired. In contrast, HSAN1-associated variants display altered substrate selectivity, shifting flux towards non-canonical 1-deoxysphingolipid (1-deoxySL) production but decreasing canonical synthesis. The variant associated with a mixed sensory-motor phenotype exhibit a third metabolic state with elevated 1-deoxySL formation and, in contrast to HSAN1-variants, increased canonical sphingolipid synthesis. Sphingolipid profiling reveals that ALS variants are characterized by preferential accumulation of dihydro- and intermediate chain sphingolipid species. Notably, the separation of lipid species between ALS and HSAN1 is robust, with canonical sphingolipids enriched in ALS variants, while long-chain 1-deoxySL dominate in HSAN1. The SPT-variants associated with mixed sensory and motor symptoms are associated with elevated levels of both types. The data support the view that segregated shifts in sphingolipid flux underlie divergence of clinical phenotypes in SPT-variants and offer guidance for therapeutic interventions. Importantly, therapeutic strategies must account for these metabolic configurations, as L-serine supplementation may benefit HSAN1 but exacerbate pathology in ALS and sensory-motor disease conditions.

## Introduction

Sphingolipids are essential bioactive lipids that are particularly abundant in neuronal tissues, where they regulate membrane architecture, cellular growth, differentiation, and signal transduction [1-3]. The rate-limiting step of *de novo* sphingolipid synthesis is catalyzed by serine-palmitoyltransferase (SPT) (Fig. 1A), which condenses palmitoyl-CoA with L-serine to generate amino alcoholic hydrocarbons called long-chain base (LCB) [4]. LCBs constitute the structural backbone of all sphingolipids and thereby define the metabolic entry point into the pathway. Mammalian SPT predominantly generates saturated C_18_-LCB called sphinganine (SA, d18:0). Following SPT-mediated synthesis of SA is N-acylated by one of six ceramide synthases (CerS1-6) to form dihydroceramides (dhCer), which are subsequently desaturated by DEGS1 to generate ceramides (Cer) (Fig. 1A). These early steps occur in the endoplasmic reticulum (ER), after which ceramides are transported to the Golgi for synthesis of complex sphingolipids, including sphingomyelin (SM) and glycosphingolipids such as hexosyleramides (HexCer). Variation in acyl chain length, saturation, and headgroup modifications generates substantial species diversity within the sphingolipid pool [5].

**Figure 1:**
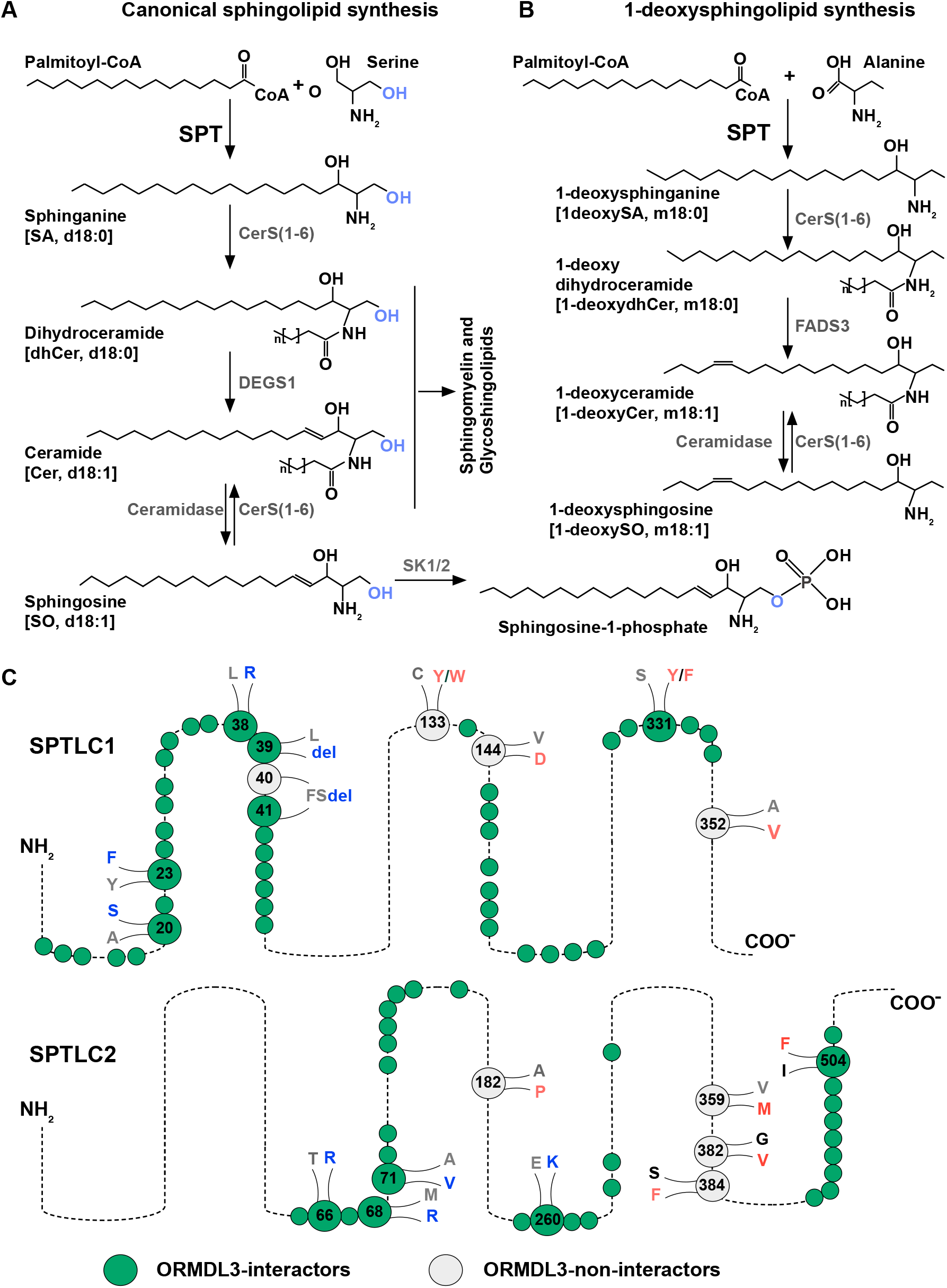
The *de novo* sphingolipid synthesis in mammals. (**A**) Canonical sphingolipid synthesis pathway. The serine-palmitoyltransferase (SPT) uses serine and palmitoyl-CoA for long-chain base synthesis. (**B**) 1-deoxySL synthesis pathway. SPT also utilizes alanine as substrate and produces 1-deoxysphingolipids that lack C_1_-OH. Key enzymes in the pathway are, CerS, Ceramide synthase; DEGS1, dihydroceramide desaturase; FADS3, fatty acid desaturase; SK1/2, sphingosine kinase 1 and 2. n is fatty acid chain length (C_16_ to C_26_). (**C**) Cartoon (not drawn to scale) depicting wild-type amino acid (grey), HSAN1 (red)- or ALS-missense and deletion (blue, del) variant positions in SPTLC1 and SPTLC2. Green circles depict ORMDL-interacting amino-acid residues. ALS mutations occur in ORMDL-interacting while HSAN1 mutations occur in ORMDL-non-interacting residues (grey circles) [44].

SPT is a multi-subunit complex in mammals. The enzymatic subunits of SPT are encoded by *SPTLC1, SPTLC2*, and *SPTLC3*. The minimal functional complex consists of a heterodimer between SPTLC1 and either SPTLC2 or SPTLC3 [6]. While SPTLC1 and SPTLC2 are ubiquitously expressed and responsible for bulk *de novo* sphingolipid synthesis, SPTLC3 is expressed in specific tissues only. Several missense mutations in *SPTLC1* and *SPTLC2* are associated with inherited sensory neuropathy type 1 (hereafter referred to as SPT-HSAN1 mutations). Belonging to Charcot-Marie-Tooth group of neuropathies, HSAN1 is an autosomal dominant neuropathy associated with loss of sensory fibers [7-10]. SPT-HSAN1 mutations shift the affinity of the enzyme from serine towards alanine, forming 1-deoxysphingolipids (1-deoxySL) (Fig 1B) [11]. 1-DeoxySL are neither converted to complex sphingolipids nor degraded, due to a missing primary hydroxyl group involved in both processes (Fig. 1A and B). Elevated 1-deoxySL levels cause cell toxicity [12]. SPTLC1 p.C133W/Y and SPTLC2 p.G382V are typical patient isolated SPT-HSAN1 variants [12-14]. 1-deoxySLs severely affect neurite formation and neuron survival in vitro [15-17], through changes in cytoskeleton and loss of actin stress fibers [18].

We and others recently characterized *SPTLC1* and *SPTLC2* variants that are associated with childhood onset juvenile ALS (hereafter referred to as SPT-ALS mutations) [19-22]. In contrast to the sensory neuropathy, all SPT-ALS patients present with characteristic motor-neuron disease phenotypes, such as spastic speech, tongue atrophy and fasciculation’s, and compromised upper and lower motor neuron function in limbs. These mutations include SPTLC1-ALS missense mutations, Y23F and L38R, as well as in frame single and double amino acid deletions of L39 and F40S41 (Fig. 1C) [19, 22]. In addition, aberrant splicing of SPTLC1 exon 2 generates variants carrying either a missense A20S substitution or complete deletion of the transmembrane domain (ex2del) [19]. The reported SPTLC2-ALS mutations are missense mutations; M68R, A71V, T166K, and E260K (Fig. 2C)[23-26].

**Figure 2:**
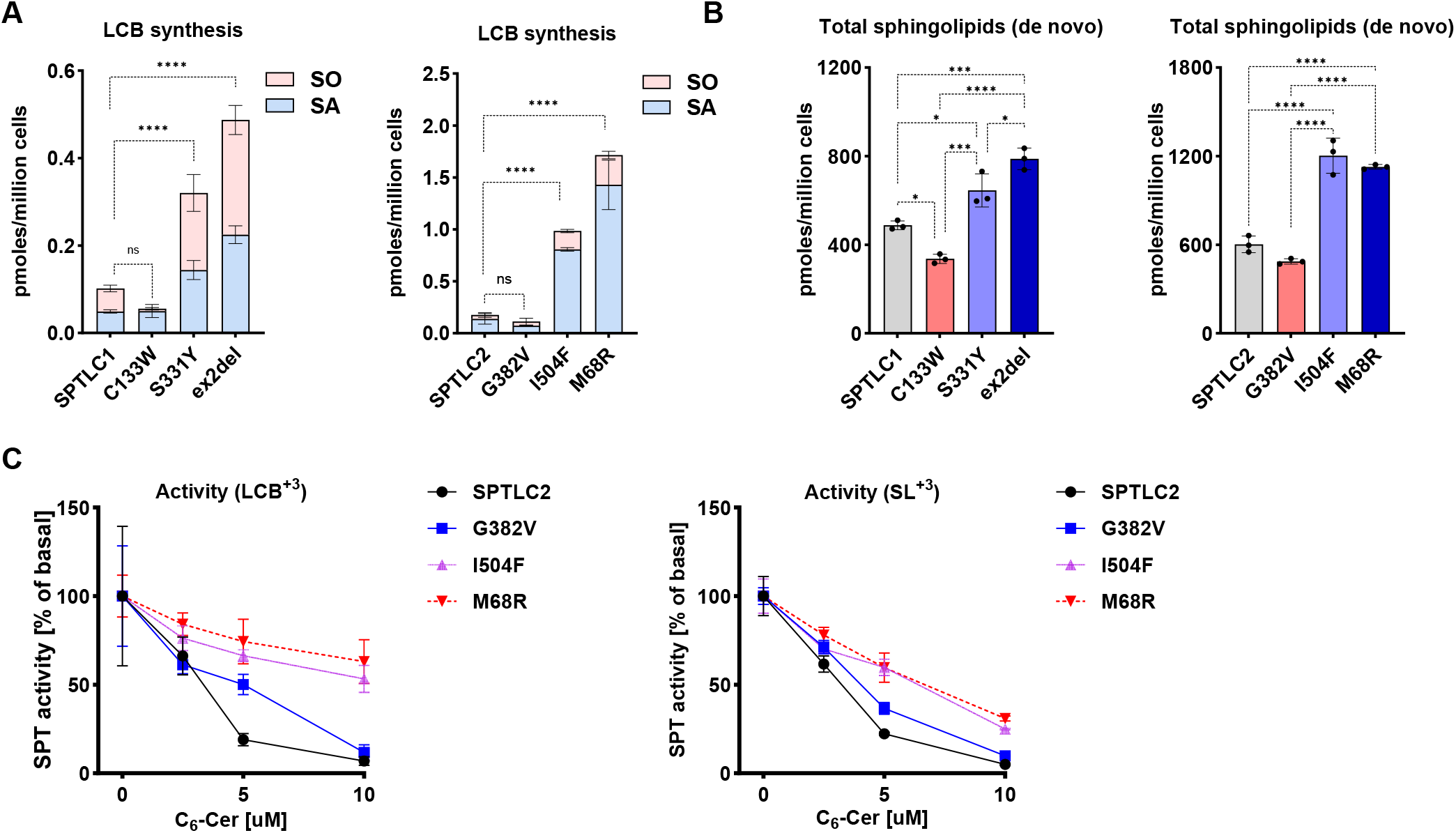
SPT activity and *de novo* sphingolipid synthesis in SPTLC1 and SPTLC2-disease variants. Levels of *de novo* produced (**A**) long chain bases (LCB), sphinganine (SA) and sphingosine (SO) as well as (**B**) total *de novo* produced sphingolipids in SPTLC1 or SPTLC2 wild-type, and corresponding variants. The *de novo* SPT activity is probed with stable isotope labelled D_3_, ^15^N-L-serine. (**C**) SPT activity in SPTLC1 and SPTLC2 wild type and indicated SPT-variants with C_6_-ceramide supplementation. Data are represented as mean ± SD, n=3 independent replicates, two-way ANOVA (for LCB plots) and one-way ANOVA with Bonferroni adjustment for multiple comparison, * p<0.05, *** p<0.001, **** p<0.0001, ns = not significant.

SPT is an important node for regulation of cellular sphingolipid levels. SPT dynamically associates with ER localized ORMDL1-3 proteins [27-29]. ORMDLs sense ceramide levels in the ER and physically interact with the SPT and markedly reducing LCB and sphingolipid synthesis. In contrast to SPT-HSAN1 variants, which are located outside ORMDL-interacting regions, SPT-ALS mutations cluster within sequence motifs that mediate interaction with ORMDL proteins [19-21, 30]. (Fig. 1C). Consistent with this, SPT-ALS patient-derived cells and plasma exhibit elevated levels of canonical sphingolipids [19-21, 23, 25, 26]

Interestingly, pathological variants such as SPTLC1 p.S331Y/F [31, 32] and SPTLC2 p.I504F [13] are associated with a relatively severe sensory phenotype as well as motor neuron dysfunction. Such patients also show muscle-wasting not observed in typical HSAN1 patients [13]. As such there have been contentions regarding the disease diagnosis of the patients.

It is not clear why distinct SPT mutations affect different neuronal populations. As the rate-limiting enzyme of sphingolipid biosynthesis, SPT functions as a metabolic pacesetter and, through its substrate promiscuity, also determines the direction of *de novo* sphingolipid production. Systemic metabolic mapping of sphingolipid alterations in SPT-HSAN1, ALS and mixed neuropathies, using controlled cellular models, is crucial to elucidate: i) sphingolipid metabolic shifts in SPT-related neuropathies, ii) correlations between lipid changes and patient phenotypes iii) sphingolipid biomarkers that may improve the diagnosis of mutation-associated diseases.

Here, we present a detailed comparison of the catalytic activities and sphingolipid species profiles of SPTLC1 and SPTLC2 variants associated with sensory, motor, and mixed sensory-motor neuropathies. By analysing how each variant alters enzyme function and reshapes the downstream sphingolipid landscape, we aim to delineate variant-specific metabolic signatures that may explain the differential vulnerability of neuronal subtypes.

## Results

### Rate of *de novo* sphingolipid synthesis in SPT mutants

We first analysed and compared the effect of representative HSAN1, ALS and sensory-motor mutations on the rate of canonical *de novo* sphingolipid synthesis. Among the SPTLC1 pathological variants, we chose C133W, ex2del, and S331Y mutations. Similarly, for SPTLC2, we chose G382V, M68R, and I504F disease variants. HEK293 T-REx cells that are SPTLC1 or SPTLC2 deficient were transfected with the wild-type or the disease variant of the corresponding gene. Inducible expression with tetracycline allows comparable expression of the constructs between lines. Presence of stable isotope labelled D_3_, ^15^N-L-serine allows *de novo* sphingolipid synthesis monitoring for comparative analysis. Incorporation of labelled amino-acid into LCB by SPT and then into downstream sphingolipids is traceable due to a mass shift (+3 Da) [20] and provides a screen shot of the rate of *de novo* sphingolipid synthesis. Expectedly, levels of *de novo* synthesized LCBs, SA and SO were elevated highest in the SPT-ALS mutants (Fig. 2A and B). These assays showed that HSAN1 mutants, SPTLC1 p.C133W and SPTLC2 p.G382V displayed lower levels of *de novo* produced free sphinganine and sphingosine as well as total sphingolipids (Fig. 2A and B).

Importantly, the sensory-motor variants SPTLC1 p.S331Y and SPTLC2 p.I504F did not mirror the reduced synthesis phenotype of HSAN1 mutants but instead exhibited synthesis that is comparable to SPT-ALS variants.

### Impaired ORMDL regulation in SPT-ALS cells

Previously, we showed that SPTLC1-ALS are refractory to ORMDL/Cer mediated inhibition while - HSAN1 mutants, such as C133W show a normal feedback inhibition[20]. To verify similar responses in SPTLC2 pathological mutants, cells were grown in presence of increasing doses of C_6_-ceramide (C_6_-Cer). As the cell-permeable ceramide analogue, C_6_-Cer inhibits LCB synthesis by sensitizing ORMDLs [33]. C_6_-Cer supplementation led to gradual dose-dependent cessation of *de novo* LCB and sphingolipid synthesis (Fig. 2C). In contrast, *de novo* activity was significantly higher in SPTLC2-ALS and sensory-motor mutants. Under these conditions, the response of HSAN1-associated G382V mutant to C_6_-Cer was similar to that of the SPTLC2 wild-type cells.

While these experiments define the rate and directionality of canonical sphingolipid flux, they do not capture potential differences in substrate utilization or species composition, which were addressed in subsequent analyses.

### Sphingolipid sub-classes in SPT-variant expressing cells

To assess flux distribution into canonical complex sphingolipids, we quantified isotope-labelled sphingolipid subclasses in mutant-expressing cells cultured in the presence of D_3_, ^15^N-serine. Because ceramide represents the central branch point of sphingolipid metabolism, we analyzed Cer, its precursor dhCer and their direct downstream products, dhSM and SM. To compare *de novo* formation of non-canonical sphingolipid formation, cells were additionally supplemented with D_4_-alanine to trace incorporation into 1-deoxySL. Like labelled serine, SPT incorporation of D_4_-alanine also adds 3 Da mass to the *de novo* generated 1-deoxySL.

Enhanced *de novo* long-chain base formation in ALS-associated SPT-variants (SPTLC1 ex2del and SPTLC2 p.M68R) results in increased incorporation into both saturated and unsaturated Cer and SM species (Fig 3A-D). In contrast, independent of the affected SPT subunit, HSAN1-associated variants accumulated 1-deoxySL (Fig. 3E and F). However, SPTLC2 p.G382V showed a more moderate increase in 1-deoxySL compared to the SPTLC1 p.C133W mutant (Fig. 3E and F), likely underscoring subunit-dependent differences in substrate promiscuity.

**Figure 3:**
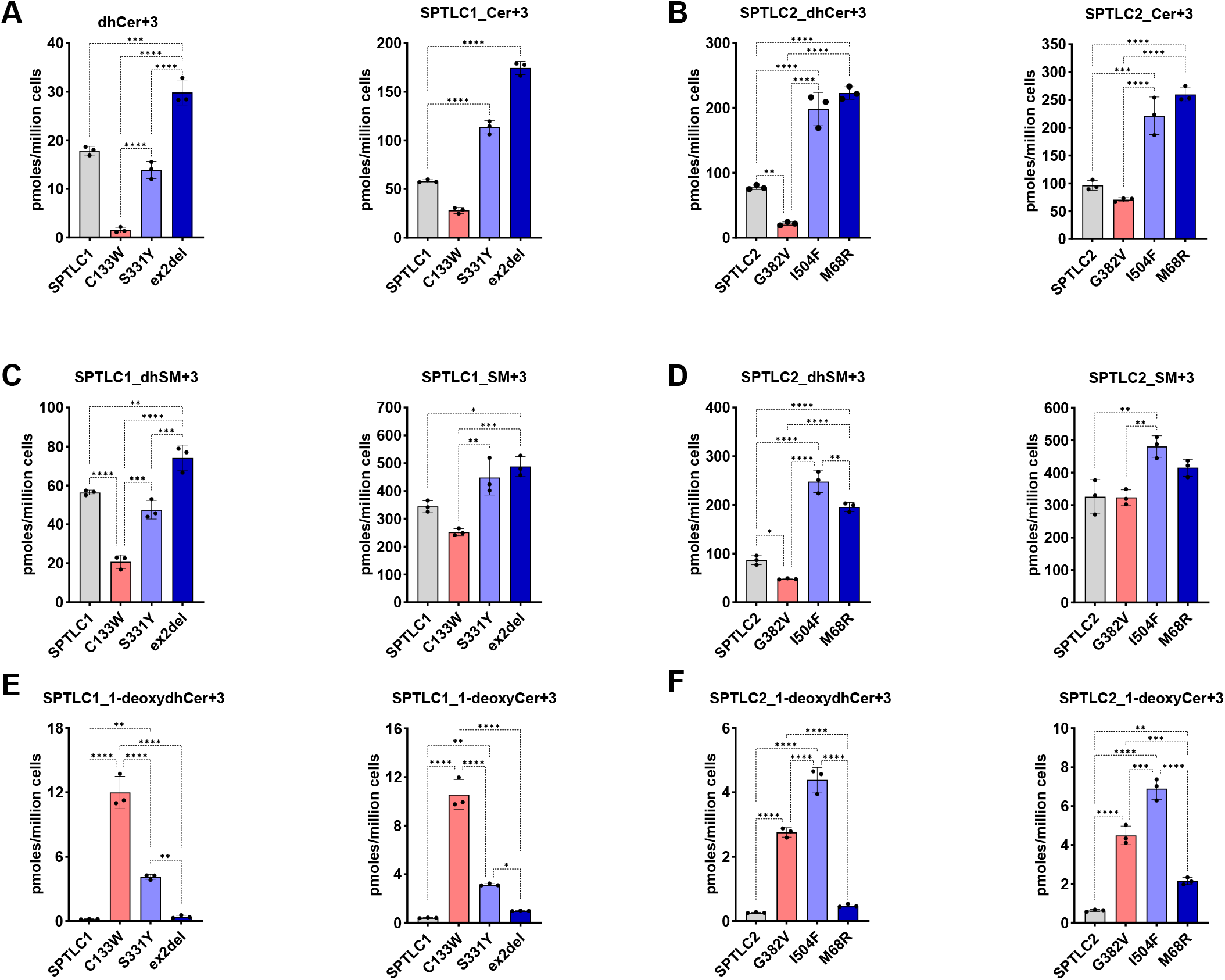
Sphingolipid sub-classes in SPT-ALS and -HSAN1 variant expressing cells. *De novo* formation of (**A-B**) dihydroceramides (dhCer) and ceramides (Cer), (**C-D**) dihydrosphingomyelin (dhSM) and sphingomyelins (SM), and (**E-F**) 1-deoxydihydroceramides (1-deoxydhCer) and 1-deoxyceramides (1-deoxyCer) in SPTLC1 and SPTLC2 knockout cells expressing the wild type and indicated variants. Cells were probed for *de novo* SPT activity by stable isotope labelling assay, with D_3_, ^15^N-L-serine and D_4_-L-alanine. Data are represented as mean ± SD, n=3 independent replicates, one-way ANOVA with Bonferroni adjustment for multiple comparison, *p<0.05, ** p<0.01, *** p<0.001, **** p<0.0001.

Sensory-motor variants defined a distinct flux configuration. In contrast to HSAN1 variants, both SPTLC1 p.S331Y and SPTLC2 p.I504F demonstrated enhanced canonical Cer and SM synthesis (Fig. 3A-D). The SPTLC1 p.S331Y variant maintained comparatively low 1-deoxySL synthesis, whereas SPTLC2 p.I504F combined increased canonical sphingolipid synthesis with robust 1-deoxySL production, exceeding that observed in SPTLC2 p.G382V (Fig. 3F).

Notably, when comparing relative fold-changes of the individual SL subclasses in mutant relative to their respective wild-type cells, dihydro-sphingolipids showed the greatest variability. While SPTLC1 p.C133W mutant showed highest decrease in all sphingolipid classes, fold-decrease in dhCer and dhSM was most pronounced in HSAN1 mutants and was independent of the affected SPT subunit (Fig. 4A-C). In contrast, ALS variants showed stronger increases in dhSM than in SM (Fig. 4C and D). Notably, SPTLC2-ALS mutants displayed more pronounced dhSL accumulation than SPTLC1-ALS mutants despite comparable increases in total complex SL levels (Fig. 4A-D).

**Figure 4:**
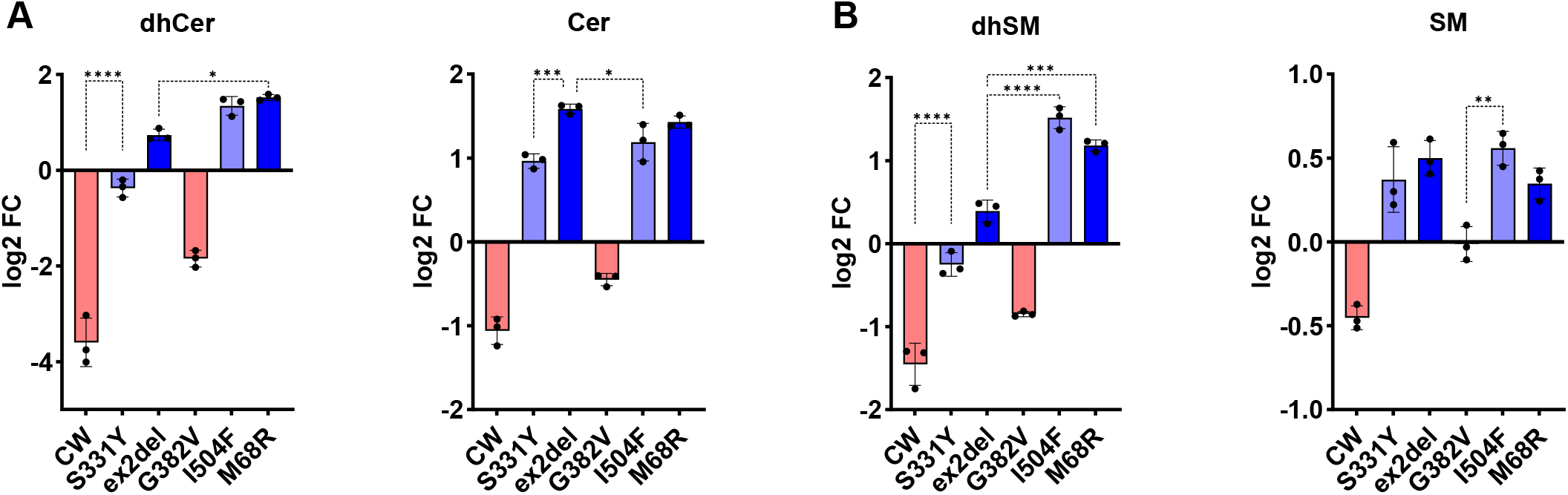
Comparison of relative changes of *de novo* produced canonical sphingolipid sub-classes. (**A**) Log_2_ fold change (log_2_FC) of dihydroceramide (dhCer) and ceramide (Cer) levels and (**B**) dihydrosphingomyelin (dhSM) and sphingomyelin (SM) levels in cells expressing disease-associated variants of SPTLC1 (C133W, S331Y, ex2del) and SPTLC2 (G382V, I504F, M68R), relative to wild-type controls. Data are presented as log_2_FCrelative to wild-type controls, shown as mean ± SD from n = 3 independent experiments. Statistical significance was determined using one-way ANOVA with Bonferroni correction for multiple comparisons, *p<0.05, ** p<0.01, *** p<0.001, **** p<0.0001.

These differences between saturated and unsaturated sphingolipid levels downstream of SPT likely reflect differential metabolic handling of dihydro- and desaturated sphingolipids rather than overall pathway flux.

Collectively, flux analysis of labelled sphingolipids delineates three metabolically distinct configurations: enhanced canonical flux with minimal non-canonical sphingolipid synthesis in ALS variants; reduced canonical flux with prominent 1-deoxySL production in HSAN1 variants; and mutation-specific composite states within sensory-motor variants characterized by elevated canonical output with variable, subunit-dependent shifts to 1-deoxySL production.

### Sphingolipid species profiles in SPT-variant expressing cells

We next sought to determine whether disease-associated variants in SPTLC1 and SPTLC2 impose discrete lipid signatures or instead distribute along a continuous sphingolipid sub-class phenotype axis. In both SPTLC1 (ex2del) and SPTLC2 (M68R) ALS-associated variants, lipid remodelling was characterized by a largely unidirectional increase in canonical sphingolipids (Fig. 5A and B). Such a shift indicates a global positive shift in canonical sphingolipid output, due to unhinged LCB synthesis of the SPT-ALS mutants. However, the most pronounced and statistically significant increase was observed in saturated and unsaturated ceramide and SM species containing C_18_, C_20_, and C_22_ acyl chains (Fig. 5A and B).

**Figure 5:**
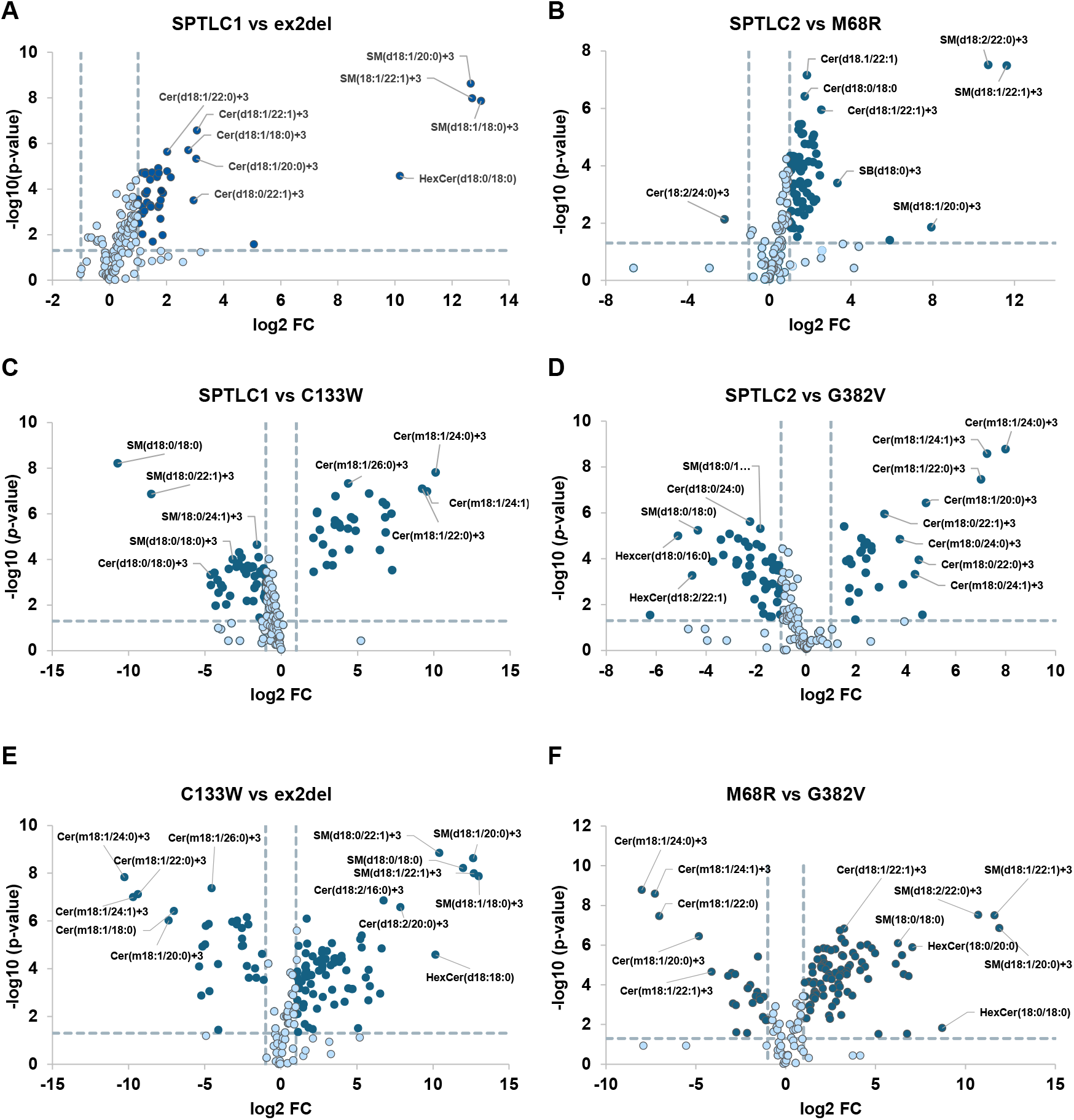
Sphingolipid species remodelling patterns associated with SPTLC1- and SPTLC2-ALS and HSAN1 disease variants. Volcano plot analysis showing changes in labelled sphingolipid species following stable isotope tracing of *de novo* sphingolipid synthesis. Selected significantly altered species are annotated. Comparison of (**A**) SPTLC1 and exon 2 deletion (ex2del) ALS variant, (**B**) SPTLC2 wild-type and the M68R-ALS variants, showing increased accumulation of canonical sphingolipids including ceramides (Cer) and sphingomyelins (SM). (**C**) Comparison of SPTLC1 and C133W-HSAN1 variant, (**D**) SPTLC2 wild type and G382-HSAN1 variant, demonstrating a shift toward increased 1-deoxySL and reduced canonical sphingolipid production. (**E**) Direct comparison of C133W and the ex2del variant, as well as (**F**) SPTLC2 M68R and G382V variants highlighting opposing sphingolipid remodelling states between HSAN1 and ALS variants. Dashed vertical lines indicate log_2_FC thresholds, and the horizontal dashed line indicates the significance threshold (p = 0.05).

In contrast, HSAN1 variants, SPTLC1 C133W and SPTLC2 G382V, displayed a reciprocal metabolic signature. These mutants showed robust accumulation of 1-deoxySL species, particularly those carrying long and very-long-chain fatty acids (C_22_-C_26_) (Fig. 5C and D). Most interestingly, multiple canonical Cer and SM species were reduced relative to wild-type controls.

To determine whether ALS- and HSAN1-associated mutations represent distinct biochemical states independent of SPT subunit context, we directly compared ALS and HSAN1 variants within each catalytic subunit. This direct comparison eliminates potential confounding effects of wild-type baseline differences observed due to intrinsic SPTLC1 and SPTLC2 activities. In line with our previous results [20], comparison of SPTLC1 C133W (HSAN1) to ex2del (ALS), showed a striking bidirectional segregation in lipid distribution (Fig 5E). Canonical ceramide and sphingomyelin species were significantly enriched in the ALS variant, whereas 1-deoxySL species were enriched in the HSAN1 variant (Fig. 5E). A nearly identical pattern emerged when comparing SPTLC2 M68R (ALS) and SPTLC2 G382V (HSAN1) (Fig. 5F). Canonical SM and Cer species with C_18_-C_22_ acyl chains were strongly enriched in the ALS variant, whereas the HSAN1 variant preferentially accumulated long and very-long-chain 1-deoxySL. The lipid species segregated robustly, clustering distinctly toward opposite sides of the volcano plot. This demonstrates that the HSAN1 and ALS mutations define clearly divergent lipid phenotypes, irrespective of the subunit.

### SPT-mixed variants lead to a distinct metabolic state

In contrast to the metabolic polarisation of ALS and HSAN1 mutations, mixed sensory-motor variants (SPTLC1 p.S331Y and SPTLC2 p.I504F) did not preferentially enrich one sphingolipid type over the other. Instead, these mutants showed concurrent and significant increases in both canonical sphingomyelin/ceramide and 1-deoxySL species (Fig. 6A and B). Importantly, the magnitude of canonical sphingolipid accumulation in these variants overlaps with that observed in ALS mutations, while the increase in 1-deoxySL overlaps with HSAN1 mutations.

**Figure 6:**
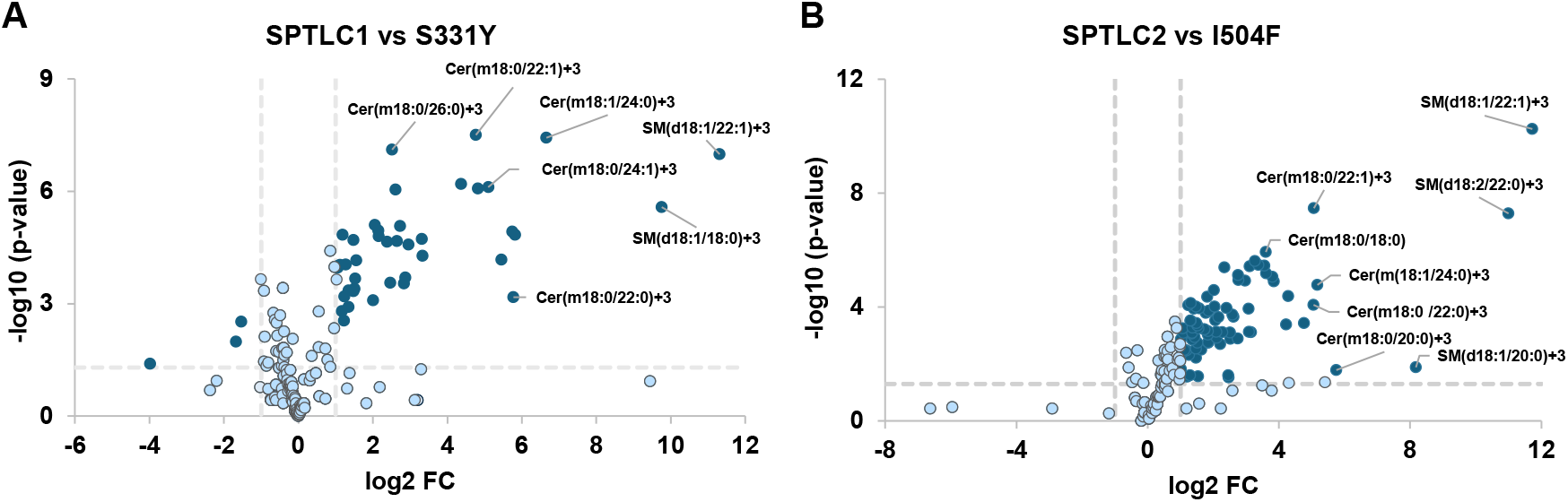
Sphingolipid species remodelling in cells expressing SPT sensory-motor variants. Volcano plots showing changes in *de novo* produced sphingolipid species following stable isotope tracing of sphingolipid synthesis. (**A**) Comparison of SPTLC1 wild-type and the sensory-motor neuropathy-associated variant S331Y, showing increased synthesis of canonical sphingolipids, particularly 1-deoxysphingolipids (m18:0 and m18:1 species) but also canonical sphingolipid such as sphingomyelins (SM). (**B**) Comparison of SPTLC2 wild-type and the sensory-motor neuropathy-associated variant I504F, demonstrating a similar increase in 1-deoxySL and labelled canonical sphingolipid species. Dashed vertical lines indicate log_2_FC thresholds, and the horizontal dashed line denotes the significance threshold (p = 0.05).

To visualize differences in sphingolipid species across variants, we performed hierarchical clustering and heatmap analysis. Lipid species segregated into two major modules corresponding to canonical sphingolipids and 1-deoxySL (Fig. 7). Canonical sphingolipids were consistently elevated in ALS-associated variants of both SPTLC1 and SPTLC2, whereas HSAN1 variants were dominated by long-chain 1-deoxySL. Compared to the respective wild types, HSAN1 mutations showed decrease in canonical lipid species. Variants associated with mixed sensory-motor phenotypes displayed a composite lipid profile with concurrent increases in both canonical and non-canonical species (Fig. 6). Together, these patterns reveal phenotype-specific sphingolipid remodelling across SPT-associated disease states.

**Figure 7:**
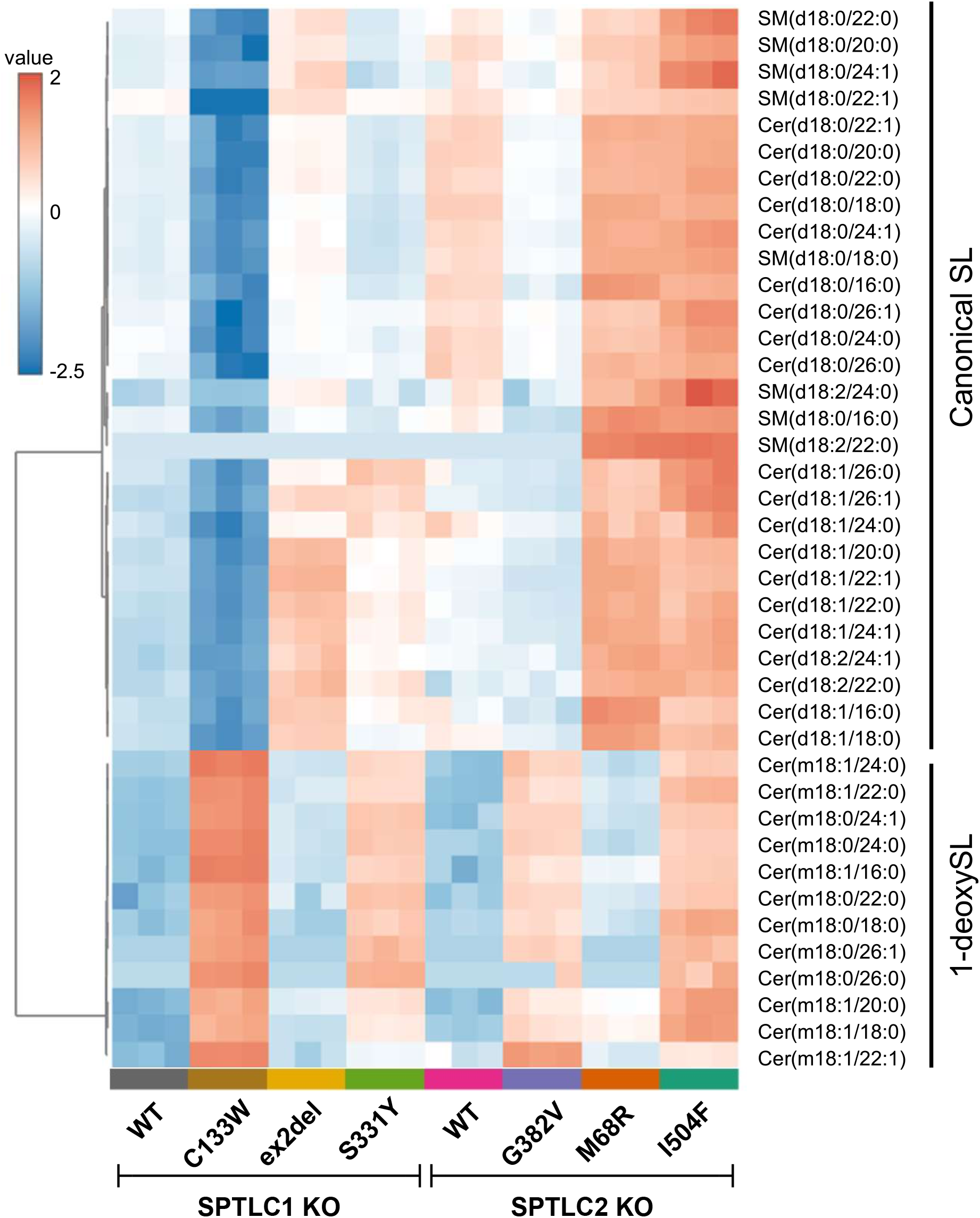
Phenotype-specific sphingolipid remodelling and segregation across SPTLC1- and SPTLC2-variants. Heatmap showing relative changes in labelled sphingolipid species following stable isotope tracing of *de novo* sphingolipid synthesis in cells expressing SPTLC1 or SPTLC2 disease variants. Columns represent the indicated genotypes grouped by gene and clinical phenotype (wild-type, HSAN1-associated, ALS-associated, and sensory-motor variants), and rows represent individual sphingolipid species. Species are separated into canonical sphingolipids (Cer and SM, d18:0 and d18:1) and non-canonical 1-deoxysphingolipids (1-deoxySL, m18:0 and m18:1). Colour intensity indicates the scaled relative abundance (z-score) of labelled sphingolipid species across samples, with red indicating increased levels and blue indicating reduced levels relative to the dataset mean. Shown is a plot of log-transformed (base 10) data with Pearson’s distance measure. The heat map was generated using MetaboAnalyst suite 6.0 [43].

## Discussion

SPT functions as the rate-limiting enzyme of sphingolipid biosynthesis and thereby establishes the metabolic set-point in neuronal sphingolipid homeostasis. In the present study, we demonstrate that disease-associated variants in both SPTLC1 and SPTLC2 define reproducible and polarized sphingolipid remodelling states that segregate according to clinical phenotype. ALS-associated variants uniformly drive enhanced canonical sphingolipid flux with impaired ceramide-mediated feedback inhibition. HSAN1-associated variants shift substrate utilization towards Ala and 1-deoxySL production at the expense of canonical output. Sensory-motor variants present a third metabolic state characterized by elevation of both canonical and non-canonical sphingolipids. Together, these findings support the concept that SPT mutations define segregated shifts in sphingolipid homeostasis relevant to each disease phenotype.

ALS mutations reside within stretches of ORMDL-interacting protein sequences of SPTLC1 and SPTLC2 (Fig. 1C). While we cannot completely rule out catalytic hyperactivity, the reduced feedback sensitivity provides a mechanistic basis for sustained canonical sphingolipid overproduction in SPT-ALS mutations (Fig. 2A-C). Importantly, this dysregulation appears to operate similarly across both SPTLC1 and SPTLC2 catalytic subunits (Fig. 2A-C), reinforcing the concept that loss of homeostatic restraint underlies the SPT-associated motor neuron disease [20]. ALS variants therefore define one extreme of the sphingolipid phenotype spectrum, marked by preferential accumulation of canonical *de novo* derived sphingolipids. In contrast, HSAN1-variants occur in non-ORMDL interacting regions of the two proteins (Fig. 1C). SPT-HSAN1 exhibit altered amino acid selectivity, leading to accumulation of 1-deoxySL and a reduced canonical output (Fig. 3A-C). Such variants mark the other extreme of the spectrum. Thus, ALS and HSAN1 arise from mechanistically distinct perturbations of SPT regulation, respectively, the feedback dysregulation and substrate shift.

Free labelled SA is the first stable product in the *de novo* pathway but represents only a fraction of the total SL pool in wild-type or SPT-mutant expressing cells. This, on one hand, shows that the flux from SPT through to the downstream enzymes is rapid and constitutive. On the other hand, a minor increase in labelled SO under these conditions argues against enhanced sphingolipid hydrolysis in ALS mutant cells (Fig. 2A and B). All SPT-ALS variants were associated with preferential accumulation of dihydroceramides and dihydrosphingomyelins, suggesting that increased SPT activity in ALS might overwhelm the capacity of dihydroceramide desaturase 1 (DEGS1), leading to an accumulation of these intermediate sphingolipids (Fig. 4). Direct comparisons between ALS and HSAN1 variants within each catalytic subunits revealed a striking bidirectional segregation of lipid species. Canonical ceramide and sphingomyelin species containing C_18_-C_22_ acyl chains are enriched in ALS variants (Fig. 5A and B), whereas long and very-long chain 1-deoxySL predominate in HSAN1 variants (Fig. 5C and D). Whether these compositional changes are limited to Cer synthase activities remains to be explored. However, the separation is robust and largely non-overlapping (Fig. 5E and F). Overall, the data suggests that LCB synthesis driven by hyperactive SPT influences species distribution or modelling along the pathway. Also, ALS and HSAN1 variants anchor opposite ends of a sphingolipid remodelling axis. Importantly, this axis is conserved for SPTLC1 and SPTLC2, indicating that the biochemical phenotype is mutation-driven rather than subunit-specific.

The neuronal subtype vulnerability may be determined by quantitative thresholds of canonical versus 1-deoxySL burden. Dihydro species, such as dhCer and dhSM influence membrane packing, bilayer rigidity, and lipid raft architecture differently from desaturated counterparts [34, 35]. Additionally, sphingolipid acyl-chain length influences membrane fluidity and is an important factor for sphingomyelin-cholesterol interaction [36, 37]. As such, sphingolipid remodelling as observed in SPT-variants may induce architectural changes in membranes that ultimately lead to cell cycle arrest, apoptosis, leukodystrophy, and neuron death [2, 38, 39]. Motor neurons, with their long axonal projections and high membrane turnover demands, may be particularly sensitive to canonical sphingolipid excess, dhSL accumulation, and altered acyl-chain composition, whereas sensory neurons are preferentially vulnerable to 1-deoxySL accumulation. The precise molecular basis of this differential sensitivity remains to be defined.

The disease phenotype correlates with the directionality of metabolic flux partitioning within the sphingolipid pathway and governed by SPT mutation type. Sensory-motor variants did not cluster exclusively with ALS or HSAN1 phenotypes but instead displayed simultaneous elevation of canonical and 1-deoxySL species (Fig. 6A-B and Fig. 7). This hybrid biochemical configuration provides a plausible explanation for mixed clinical presentations. The coexistence of increased canonical sphingolipids and metabolically-resistant 1-deoxySL may impose compounded metabolic stress on both motor and sensory neurons. However, the magnitude and consistency of flux enrichment across both SPTLC1 and SPTLC2 sensory-motor mutations suggest that these variants define a reproducible metabolic configuration distinct from both the low-flux and 1-deoxySL generating HSAN1 state and the canonical ALS pattern. The remodelling pattern showed that sensory-motor variants in SPTLC1 and SPTLC2 do not represent partial shifts along a continuum as suggested earlier [40] but instead induce a combined activation of canonical and atypical sphingolipid pathways.

The mechanistic contradiction between substrate shift (HSAN1) and feedback dysregulation (ALS) has direct therapeutic relevance. L-serine supplementation ameliorates 1-deoxySL accumulation in HSAN1 [15, 41] but would be predicted to exacerbate canonical sphingolipid overproduction in SPT-ALS or sensory-motor variants. [19, 20, 40]. In addition to this, inhibiting synthesis of long-chain fatty acids (C_24_, C_26_) pharmacologically also rescues 1-deoxySL induced neuron death [17]. This approach may prove counter-productive in ALS and sensory-motor neurons as the short and mid-chain length canonical sphingolipids are expected to increase under this condition (C_16_-C_22_). Thus, therapeutic strategies must be tailored to the underlying metabolic configuration rather than the affected gene alone.

### Limitations of the study

This study relies primarily on overexpression systems in SPT-deficient HEK293 cell models, which may not fully recapitulate neuronal context. Spatial lipid distribution, compartment-specific flux, and long-term adaptive responses were not assessed. Future work in motor and sensory neuron-derived models and *in vivo* systems will be required to determine how these metabolic alterations translate into selective neuronal degeneration.

## Materials and Methods

### Cell culture

Flp-In T-REx cells were cultured in high-glucose Dulbecco’s Medium (DMEM, Sigma-Aldrich, St. Louis, MO, USA) supplemented with 10% fetal bovine serum (FBS) and 1% Penicillin/Streptomycin (P/S). Cells were incubated at 37°C in a humified atmosphere by 5% CO2. The establishment of the SPTLC1-knockout (KO) and the SPTLC2-KO Flp-In T-REx cell lines and generation of SPT-mutant expressing cells have been described previously [20, 42]. All plasmid constructs used in the experiments were generated following standard molecular biology protocols. Transfections of plasmids into HEK293 cells were carried out using Lipofectamine 3000 (Thermo Fisher Scientific, MO, USA).

### Protein expression in KO cells

Plasmids containing FLAG-tagged SPTLC1 and SPTLC2 wild-type or pathogenic variants were transfected in the respective KO cells. For this, cells were plated in 6 well plates (200, 000 cells per ml) in DMEM media without antibiotics. After 24 hours, 2 µg of plasmid DNA was transfected. After 24 hours of transfection, protein expression was induced with 1 μg/mL tetracycline for 4 hours in DMEM (Thermo Fischer Scientific).

### Stable isotope labelling assay in cells

Sphingolipid labeling assays and serine-palmitoyltransferase (SPT) activity measurements were performed using transiently transfected cells. For standard labeling assays, the medium was replaced with L-serine-free DMEM (Genaxxon Bioscience) supplemented with 10% FBS and 1 mM D_3_-^15^N-L-serine and 2 mM D_4_-L-alanine (Cambridge Isotope Laboratories). The cells were incubated in this labeling medium for 16 hours. When included, C_6_-ceramide (C_6_-Cer) was added concurrently with the labeling medium for activity assays. The cells were harvested in ice-cold PBS. 50 μl aliquot was taken for cell counting using a Z2 Coulter Counter (Beckman Coulter), and cell pellets were stored at -20 °C until lipid extraction.

### Lipid extraction and analysis

Sphingolipidomic analysis was essentially conducted as described previously[20]. Briefly, frozen cell pellets were resuspended in 50 μl PBS and extracted with 1 ml of MMC lipid extraction solution [methanol/MTBE/chloroform (4:3:3, v/v/v)]. Extractions were performed in 2 ml safe-lock Eppendorf tubes at 37 °C using a thermomixer (1400 rpm, 60 min). The resulting single-phase supernatant was collected, dried under nitrogen, and reconstituted in 100 μl methanol. Lipid separation was carried out on a C18-Waters LC column (150 mm × 2.1 mm, 2.6 μm particle size) using a Transcend UHPLC system (Thermo Fisher Scientific). Details of Liquid chromatography used and methods for identification and quantification of lipids are described here [20].

### Statistics

Unless otherwise specified, data were analyzed using one-way ANOVA followed by Bonferroni’s correction for multiple comparisons. xx were analyzed using two-way ANOVA with Dunnett’s post hoc test. All results are presented as mean ± standard deviation (SD). Adjusted P values less than 0.05 were considered statistically significant. Statistical analyses were conducted using GraphPad Prism version 10.0 (GraphPad Software, Inc.) and MetaboAnalyst version 6.0 [43].

## Author contributions

MAL designed and performed experiments. NZ performed experiments, analyzed and interpreted data. NZ, TH and MAL wrote and reviewed the manuscript. All authors approved the manuscript.

## Acknowledgements

This work was supported by grants from *Schweiz Stiftung für die Erforschung der Muskelkrankheiten* (FSRMM) and the EMPIRIS foundation to MAL and by the Swiss National Science Foundation (SNF 310030_215134) and by the SNF under the frame of the European Joint Program on Rare Diseases (EJP RD+SNF 32ER30_187505) to TH.

